# Therapeutic targeting of KRAS mutation-driven tumorigenesis by extracellular vesicles loaded with small interfering RNA

**DOI:** 10.1101/2024.01.17.576015

**Authors:** Yanhua Zhai, Shuiqin Niu, Tao Qiu, Yuan Yi, Yu Yan, Rui Hu, Xinjun He, Ke Xu

**Affiliations:** Vesicure Therapeutics Co. Ltd., Biobay 3B, Suzhou Industrial Park, Suzhou, 215000, China

## Abstract

*RAS* is a well-known oncogene contributing to significant proportion of cancer incidences, yet it remains as an undruggable target to majority of therapeutic modalities. Here, we explored the potential of extracellular vesicles (EVs) for therapeutic targeting of *KRAS* mutation-driven tumorigenesis. EVs loaded with small interfering RNA (siRNA) against *KRAS* successfully inhibited tumor xenograft growth when injected intratumorally in mice models. Intriguingly, when injected intravenously, EVs were still able to accumulate in tumors and deliver *KRAS* targeting siRNA payload in sufficient amount to show tumor growth inhibition. Therefore, EV-siRNA platform show great promises for therapeutic targeting of *KRAS* mutations and other undruggable targets.

## Introduction

Mutations in the *RAS* oncogenes are well-known genetic drivers for numerous cancer types, including non-small cell lung cancer (NSCLC), colorectal cancer (CRC), and pancreatic ductal adenocarcinoma (PDAC) (Huang et al. 2021; Zhu et al. 2022). Specifically, *KRAS* mutations are heavily concentrated in hotspot codons such as G12, G13 and Q61, rendering KRAS protein in abnormally activated state (Huang et al. 2021; Zhu et al. 2022). While *KRAS* mutants have been primary drug targets in anti-cancer therapy, it has been considered to be undruggable despite tremendous efforts with small molecule inhibitor development. Until recently, the breakthrough was made by AMG510 (Sotorasib) and MRTX849 (Adagrasib) targeting the G12C mutation, showing marked clinical responses across multiple tumor types (Blair 2021; Ou et al. 2022). Nonetheless, a larger portion of *KRAS* mutation types are still out of reach without approved therapeutic options.

Small interfering RNA (siRNA) therapeutics is emerging as an innovative modality of medicines, providing new possibilities to yet-to-overcome diseases (Hu et al. 2020). Mechanistically, siRNA pair with complementary mRNA leading to target degradation, thereby potently silencing gene expression. There has been significant progressive with development of siRNA drugs including Nedosiran, Inclisiran, Vutrisiran, Lumasiran, Givosiran, Patisiran etc. In theory, siRNA can be rationally designed to silence virtually any genes, which is highly advantageous for these considered undruggable targets by small molecules or antibodies.

Extracellular vesicles (EVs) are nano-sized, membrane-bound vesicles secreted by most cell types, which hold great promises as delivery platform for various therapeutics (Kowal, Tkach, and Thery 2014; Vader et al. 2016). Previously we have successfully engineered EVs as efficient platform for functional siRNA delivery both *in vitro* and *in vivo* (Qiu et al. 2023). Here, we explored the possibility of utilizing EVs to deliver siRNA to target *KRAS* mutation-driven tumorigenesis.

## Results

### EV-siR-*KRAS* efficiently inhibited tumor xenograft growth through intratumoral injections

Previously we have achieved high density siRNA loading into EVs lumen, through home-developed dilEVry™ loading system (Qiu et al. 2023). Here as proof-of-concept study, EVs loaded with *KRAS* mutation-targeting siRNA were tested for its ability to inhibit tumor growth *in vivo*. Lung carcinoma cell lines A549 bearing KRAS G12S mutation and NCI-H358 bearing KRAS G12C were used to establish *KRAS*-mutation driven tumorigenesis xenograft model in nude mice. First, EV-siR-*KRAS* were intratumorally injected for eight consecutive days in A549 xenograft mice model, followed by tumor growth monitoring for another two weeks post-treatment (**Figure 1A**). In this test, we observed that the A549 tumor xenograft expanded rapidly in control group (**Figure 1B**). In contrast, EV-siR-*KRAS* group significantly inhibited tumor growth as measured by tumor volume and the relative rate of tumor proliferation (T_RTV_/C_RTV_) (**Figure 1B-C**). Moreover, EV-siR-*KRAS* were safe and well-tolerated by the animals (**Figure 1D**). Therefore, we validated that EV-siR-*KRAS* is efficient in inhibiting *KRAS* driven tumor growth in mice model *in vivo*.

**Figure 1:**
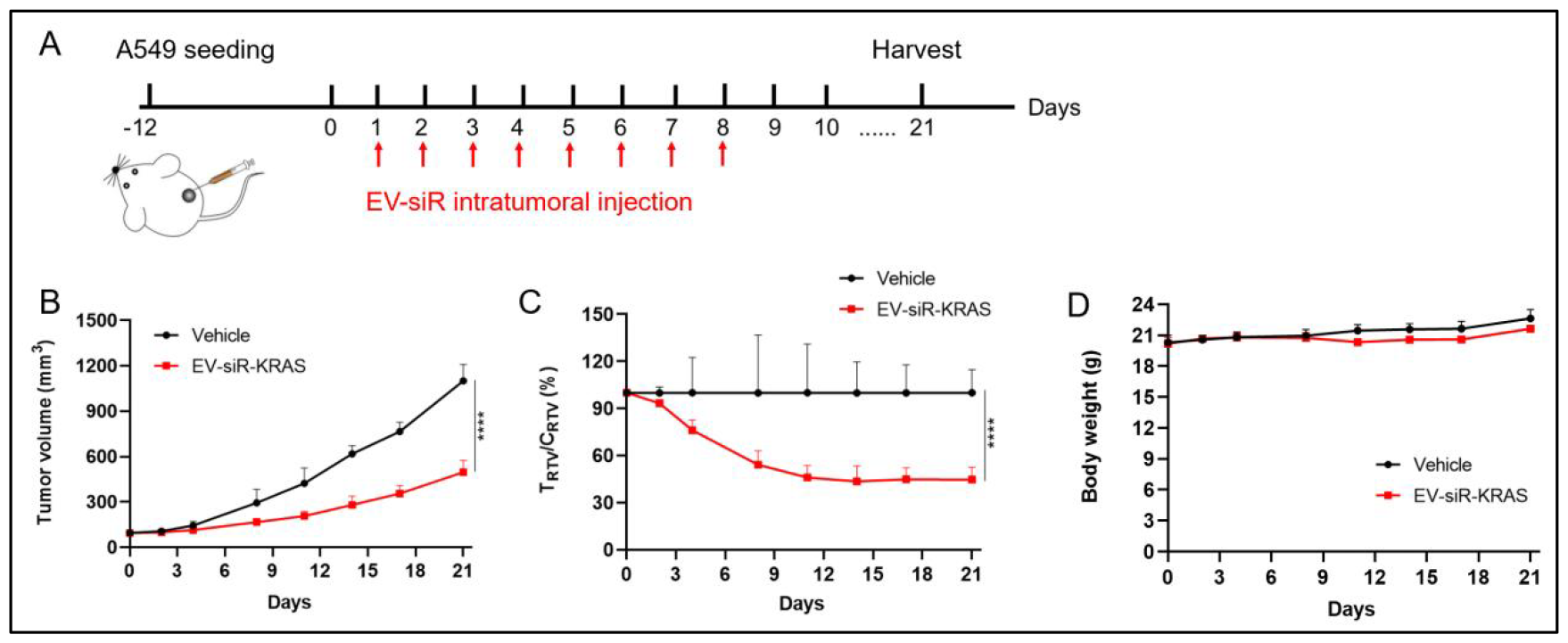
EV-siR-*KRAS* efficiently inhibited tumor xenograft growth through intratumoral injections. (**A**) Schematic of A549 tumor xenograft model in nude mice. (**B**) Tumor volume tracking for mice receiving different treatment groups. (**C**) Relative tumor proliferation rate tracking for mice receiving different treatment groups. (**D**) Body weight of animals receiving different treatment groups. ^****^: p<0.0001.

### EV-siR-KRAS efficiently inhibited tumor xenograft growth through intravenous injections

To expand the applicability of EV-siR-*KRAS* platform, we set out to explore if systemically injected EVs were still capable of accumulation and functional delivery to the tumor site. Intriguingly, we observed significant EVs accumulation in the tumor following intravenous (I.V.) injection (**Figure 2A**). Next, a G12C mutation harboring NCI-H358 tumor xenograft model was again established and EVs loaded with siRNA targeting the G12C mutation were injected intravenously (**Figure 2B**). As shown in Figure 2C-D, EV-siR-*KRAS* were able to significantly inhibit tumor growth as determined by tumor volume and the relative rate of tumor proliferation (T_RTV_/C_RTV_). Finally, we found again that EV-siR applied systemically were safe and well-tolerated by the animals (**Figure 2E**).

**Figure 2:**
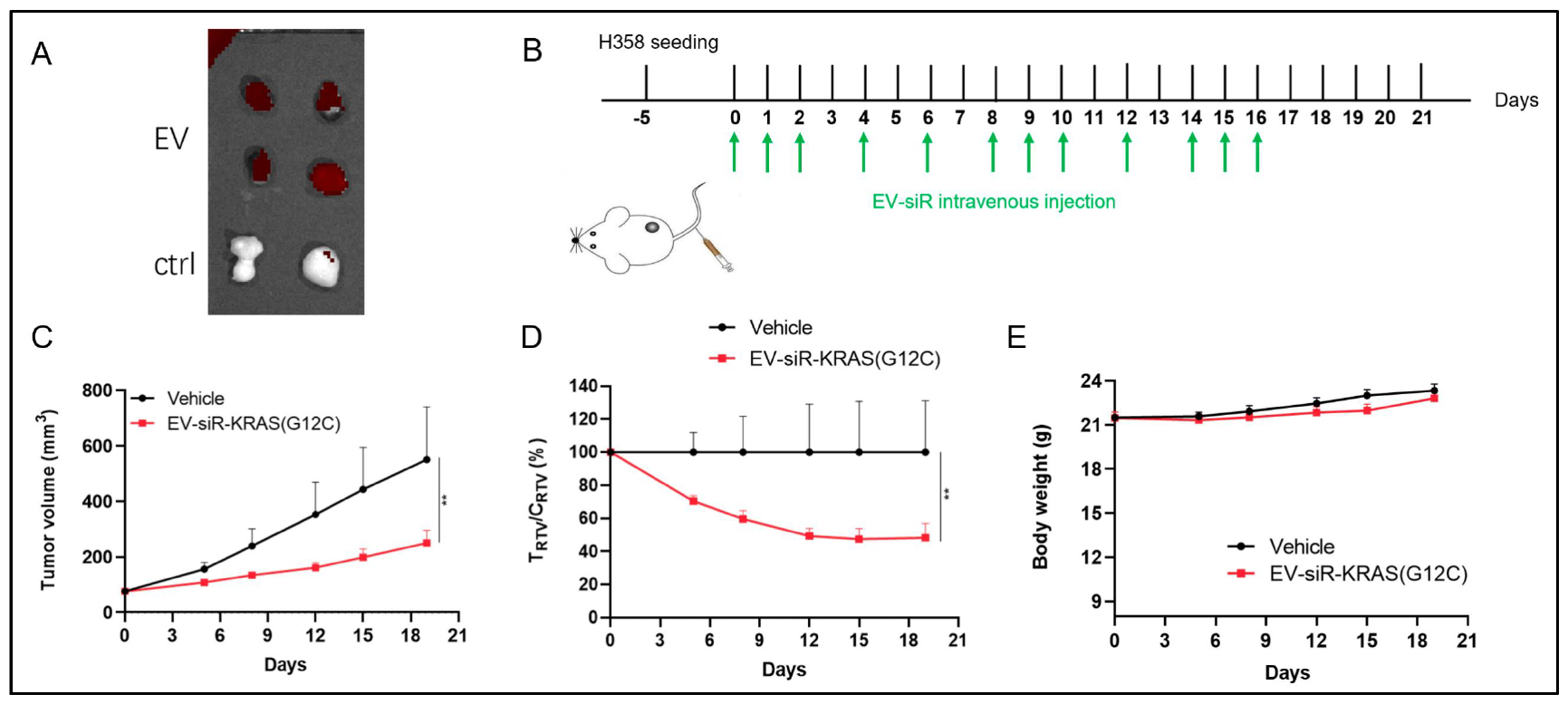
EV-siR-*KRAS* efficiently inhibited NCI-H358 tumor xenograft growth through intravenous injections. (**A**) Tumor imaging showing DiR-labeled EV accumulation in tumors from animals receiving EVs intravenously versus control group. (**B**) Schematic of H358 tumor xenograft model in nude mice. (**C**) Tumor volume tracking for mice receiving different treatment groups. (**D**) Relative tumor proliferation rate tracking for mice receiving different treatment groups. (**E**) Body weight of animals receiving different treatment groups. ^**^: p<0.01.

In summary, EV-siR-*KRAS* demonstrated efficacy in various tumor xenograft model driven by different *KRAS* mutations.

## Discussion

In our previous report, we have engineered extracellular vesicles as efficient siRNA delivery tool (Qiu et al. 2023). Here, we made explorations to expand the applicability of this platform further. EVs loaded with siRNA targeting *KRAS* oncogene successfully inhibited tumor growth driven by *KRAS* mutation, suggesting the possibility to apply this delivery platform for targeting many more “undruggable” targets. Importantly, tumorigenesis driven by distinct *KRAS* mutations can be efficiently targeted by simply changing the siRNA sequences, while keeping the same EVs delivery platform. This is highly advantageous compared to small molecules or antibody therapeutics, which all require *de novo* design for each new mutant. Finally, EVs applied through system injections were still able to accumulate to tumor site and showed therapeutic efficacy, suggesting exciting possibility of EV platform with tumor targeting ability *in vivo*. We will continue to explore, and aim for ultimately translating EVs into therapeutics.

## Materials and Methods

### Cell culture

A549 cells were purchased from Procell, and cultured in Ham’s F-12K medium (Gibco) supplemented with 10% fetal bovine serum (Gibco) and 1 × penicillin-streptomycin (Gibco). NCI-H358 cells were purchased from Procell, and were cultured in RPMI1640 medium (Gibco) supplemented with 10% feta bovine serum (Gibco) and 1 × penicillin-streptomycin (Gibco). Adherent cell culture were maintained in a humidified incubator at 37°C with 5% CO2.

### Purification of extracellular vesicles (EVs)

The HEK293 cell culture supernatant was first centrifuged at 133,900 g at 4 °C for 60 min. The crude EVs pellet were resuspended in PBS, and further layered onto 17.5% iodixanol/45% iodixanol gradient, followed by centrifugation at 150,000 g at 4 °C for 16 h. The extracellular vesicles appeared as a white layer between PBS/17.5% iodixanol, which were carefully pipetted out and washed with PBS by centrifugation at 135,000 g at 4 °C for 3 h. The refined EVs were finally resuspended in PBS.

### Tumor xenograft model

Female, 8-weeks-old nude mice (Vital River) were implanted with 1E6 cells/mice under the right fat pad region. When the average tumor volume reached around 60-80 mm^3^, the mice were randomly grouped for different treatment conditions. Intratumoral /intravenous injections and tumor volume measurement were performed according to the scheme. On last day, mice were sacrificed and tumors were excised out and imaged.

